# Germline *de novo* mutation clusters arise during oocyte aging in genomic regions with increased double-strand break incidence

**DOI:** 10.1101/140111

**Authors:** Jakob M. Goldmann, Vladimir B. Seplyarskiy, Wendy S.W. Wong, Thierry Vilboux, Dale L. Bodian, Benjamin D. Solomon, Joris A. Veltman, John F. Deeken, Christian Gilissen, John E. Niederhuber

## Abstract

Clustering of mutations has been found both in somatic mutations from cancer genomes and in germline *de novo* mutations (DNMs). We identified 1,755 clustered DNMs (cDNMs) within whole-genome sequencing data from 1,291 parent-offspring trios and investigated the underlying mutational mechanisms. We found that the number of clusters on the maternal allele was positively correlated with maternal age and that these consist of more individual mutations with larger intra-mutational distances compared to paternal clusters. More than 50% of maternal clusters were located on chromosomes 8, 9 and 16, in regions with an overall increased maternal mutation rate. Maternal clusters in these regions showed a distinct mutation signature characterized by C>G mutations. Finally, we found that maternal clusters associate with processes involving double-stranded-breaks (DSBs) such as meiotic gene conversions and *de novo* deletions events. These findings suggest accumulation of DSB-induced mutations throughout oocyte aging as an underlying mechanism leading to maternal mutation clusters.

---

*De novo* mutations (DNMs) arise spontaneously in parental gametes and result in approximately 50-100 germline mutations in their offspring^1-4^. As such, DNMs are both drivers of evolution as well as a common cause of sporadic disorders. The exact number of DNMs is highly correlated with paternal age and, to a lesser extent, with maternal age^2-4^. The paternal age effect, giving rise to about one additional DNM in the offspring per year of life of the father before conception, is thought to be due to the higher number of cell divisions that sperm cells of older men have undergone prior to this period. The mechanisms underlying the maternal age effect, giving rise to about one additional DNM per 4 year of life of the mother, are still unknown. Approximately 2-3% of all DNMs in an offspring occur in close spatial proximities (below 20kb) as clustered mutations^4-9^. These clustered DNMs (cDNMs) have a distinct nucleotide substitution spectrum with an enrichment of OG mutations, suggesting mutational mechanisms different from unclustered DNMs^4,7,8,10^. The precise composition of the mutation spectrum also varies with the inter-mutational distances of the clusters^8,11^. Contrary to unclustered DNMs, no paternal bias has been observed for the number of cDNMs^7,10^. Here, we investigated cDNMs, their potential contribution to the paternal and maternal age effect on the total number of DNMs, and the possible mechanisms underlying their occurrence.

Whole genomes of 1,291 parent-offspring trios from the Inova Translational Medicine Institute longitudinal childhood study cohort were sequenced using Illumina HiSeq2000 with average 40x coverage by Illumina services (La Jolla, USA; **Table 1, Supplementary Table 1).** This cohort was not selected for health or disease, and represents a sample of the general population delivering at a single hospital^12^. After quality control, we identified 73,755 high-confidence DNMs using a random forest classifier **(Supplementary Methods, Supplementary Table 2).** We defined cDNMs as DNMs within the same individual with all pair-wise inter-mutational distances smaller than 20kb. In total we identified 1,796 cDNMs (2.4% of all DNMs) distributed across 799 clusters, with 2-10 mutations per cluster of which 678 clusters (85%) consisted of exactly two mutations **(Supplementary Tables 3-6).** By performing read-phasing, we successfully identified the parent-of-origin for 700 cDNMs (39.0% of all cDNMs) across 400 clusters **(Table 1, Supplementary Table 7).** In line with our expectations, in 98.0% (196/200) of the fully phased clusters, all cDNMs arose on the same allele. In contrast to unclustered DNMs, we did not observe an excess of cDNMs on the paternal allele (108 maternal clusters and 88 paternal clusters, chi-square goodness-of-fit p=0.15). In addition, we created a validation dataset based on four independent studies with phased DNMs from whole-genome sequencing (WGS)^4,7,8,10^, resulting in a total of 1,643 cDNMs across 745 clusters **(Table 1, Supplementary Table 8).**

**Table 1:** Overview of cohorts

| Cohort |  | Total number | Paternal number | Maternal number |
| --- | --- | --- | --- | --- |
| Primary cohort | Probands | 1,291 |  |  |
|  | DNMs | 73,755 | 20,196 | 5,547 |
|  | cDNMs | 1,796 | 323 | 377 |
|  | Clusters | 799 | 110 (+88) | 94 (+108) |
| Replication cohort | Probands | 1,557 |  |  |
|  | DNMs | 74,395 | 9,466 | 2,796 |
|  | cDNMs | 1,643 | 133 | 195 |
|  | Clusters | 745 | 40 (+49) | 67 (+46) |

Numbers of probands, DNMs, cDNMs and clusters of the cohorts used in this study. The numbers in brackets indicate clusters where not all cDNMs could be phased for the respective parent.

To investigate the contribution of cDNMs to the parental age effects, we used a linear regression model to correlate the age of each parent with the number of phased cDNMs in the offspring. Although the number of paternal cDNMs did not show a significant correlation with the paternal age (p=0.087), we found a highly significant correlation of maternal cDNMs with maternal age (p<10^-10^), accounting for 23% (c.i. 7-38%) of the maternal age effect **(Supplementary Figures 1-2).** This effect was similar in our replication cohort (p=0.0159 and p=0.317 respectively) albeit not reaching statistical significance **(Supplementary Figure 3).** While in the primary cohort, only 5% of the probands with the youngest mothers had one or more maternal cDNMs per genome, this was more than 5 times higher (risk ratio test, p=l.4×10-11; c.i. 3.0-9.4) in probands from the oldest mothers (27% having a maternal cDNM, **Figure la).** This difference was not significant for the paternal cDNMs (13% vs 19%; risk ratio test p=0.08; 95% c.i. 0.95-2.12). In the replication cohort, the risk ratio was 2.7 for maternal cDNMS (c.i. 1.08-6.73; p=0.025) and 0.93 (c.i. 0.41-2.13; p=0.867) for paternal cDNMs.

**Figure 1:**
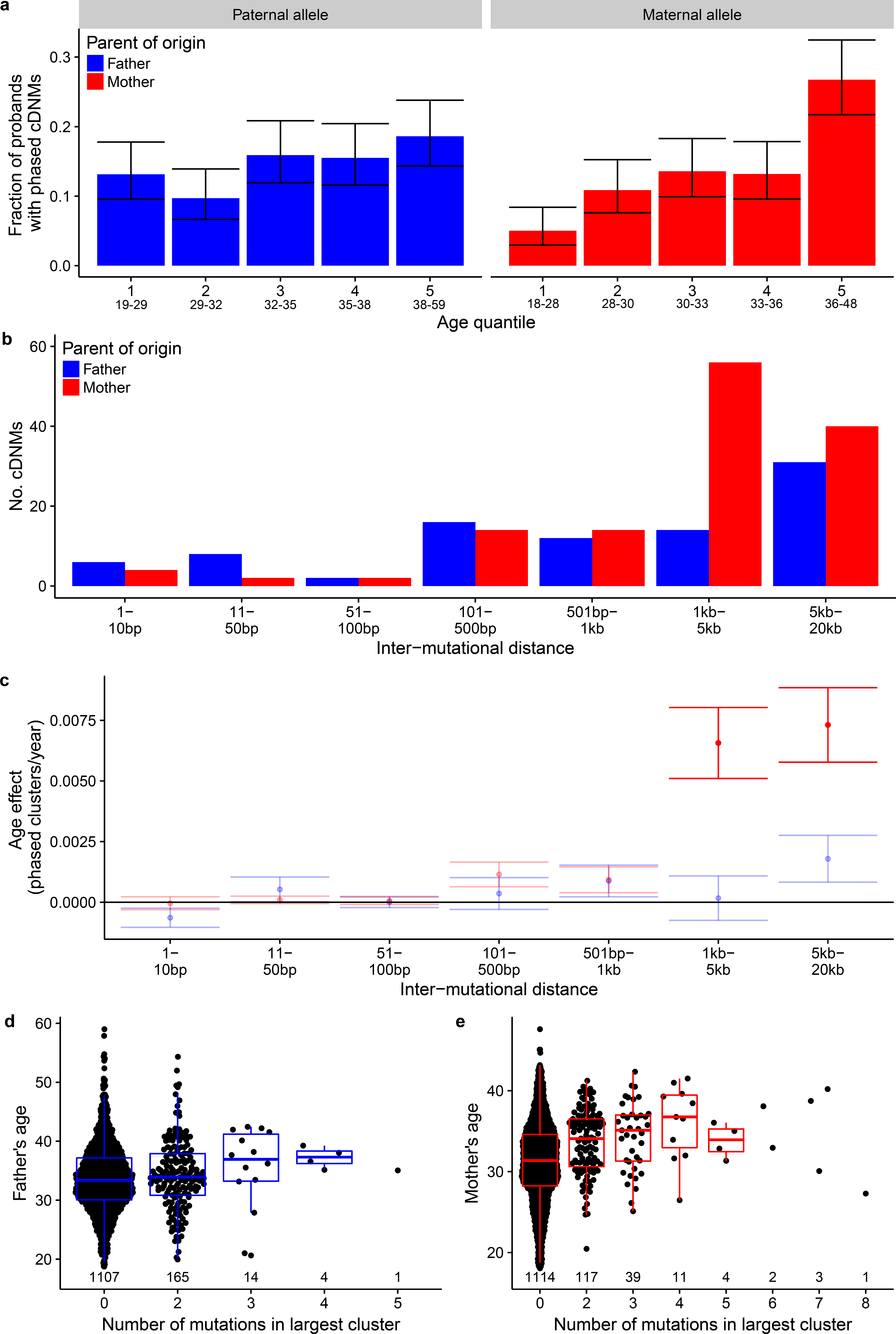
Differences between maternal and paternal cDNMs **(a)** The number of paternal and maternal cDNMs (y-axis) stratified by the distance to the nearest other cDNM (x-axis). **(b)** The fraction of probands with maternal and paternal clustered mutations (y-axis), grouped by parental age quantiles. Error bars indicate the binomial 95% confidence intervals. Labels on the lower axis indicate age ranges of the respective groups. See **Supplementary Figure 2** for graphs and regression lines, **(c)** The size of paternal and maternal age effect (y-axis) by inter-mutational distance (x-axis). Whiskers indicate the 95% confidence interval, **(d)** Age of fathers at conception **(e)** and age of the mothers at conception (y-axis) by the number of mutations in the offspring’s largest mutation cluster originating from the respective parent (x-axis). Numbers indicate the size of each group. Boxplot compartments: box: interquartile range; line: median; whiskers: extreme values <1.5 x interquartile ranges from box borders).

Previous studies showed differences in mutation profiles for clusters with different inter-mutational distances^8,11^. We found that this maternal age effect of clusters stems mostly from clusters with intra-mutational distances greater than 1kb **(Figures lb,c, Supplementary Tables 9 and 10).** Strikingly, the maximum number of DNMs in the clusters of an individual, correlates positively with maternal age (p<10^-10^, replication cohort p=0.001), but only marginally significant with paternal age (p=0.050, replication cohort p=0.829, **Figure 1d,e).** These results show that maternal clusters contain more cDNMs with larger inter-mutational distances.

We previously observed that maternal DNMs are enriched within specific genomic regions on chromosomes 8 and 16^4^. In this study, we found that 58.4% of maternal cDNMs localize to chromosomes 8, 9 and 16 (p<10^-16^, replication cohort p<10^-16^, Chi-square test; **Figure 2a, Supplementary Figures 4 and 5).** This in contrast to paternal cDNMs for which the number correlates with chromosome length (R^2^=0.72, p=6*10^-7^, replication cohort R^2^=0.25, p=0.18). The maternal cDNMs on these three chromosomes occur specifically in regions that are also enriched for maternal unclustered DNMs **(Figure 2b, Supplementary Figures 6 and 7)** and their mutation spectrum is strongly enriched for C>G substitutions compared to other maternal cDNMs **(Figure 2c,d,** bootstrapping p=0.022). This suggests a different mutational mechanism for maternal cDNMs in these regions compared to the rest of the genome.

**Figure 2:**
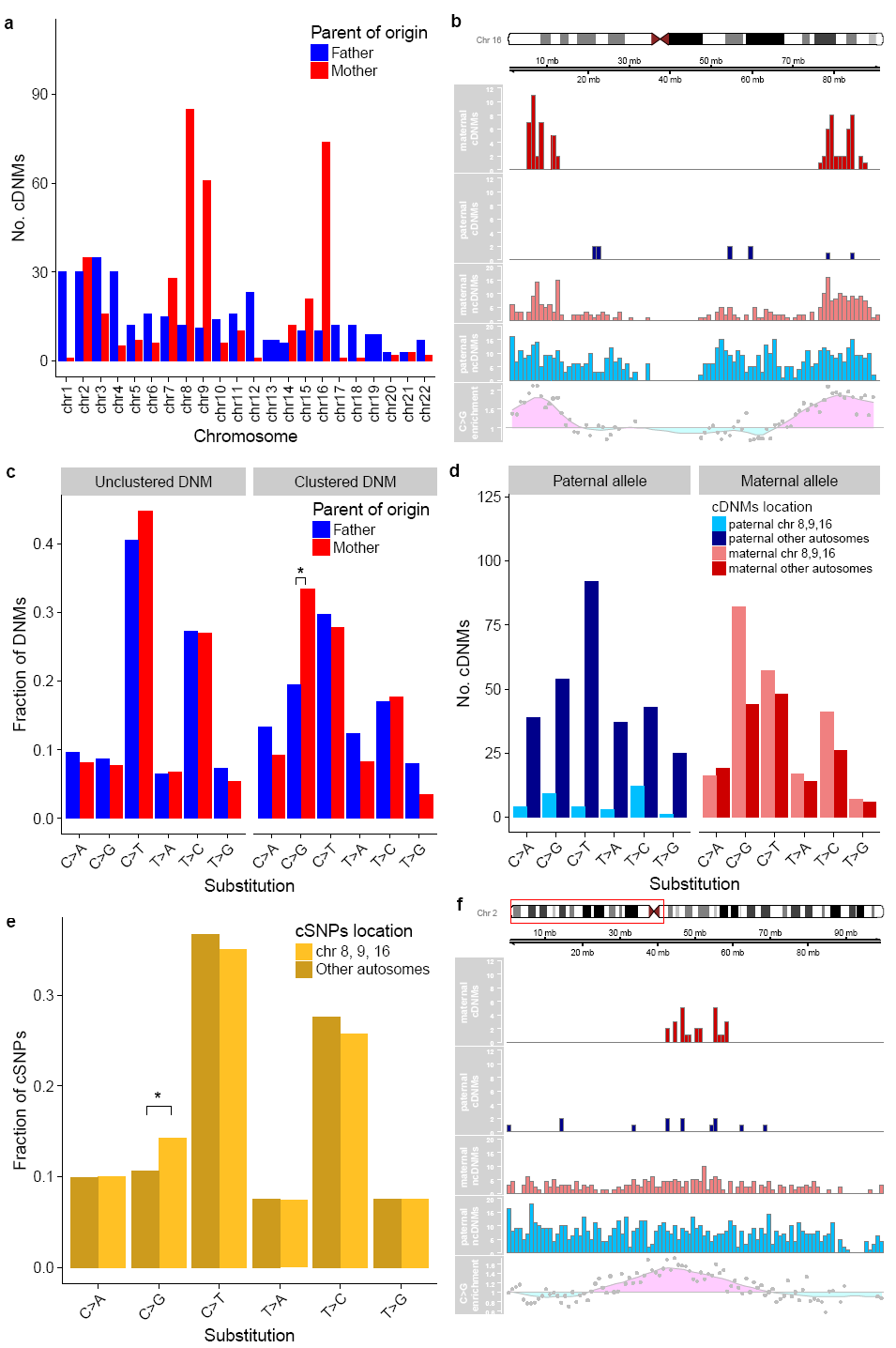
Patterns of cDNMs across the chromosomes, **(a)** The number of phased cDNMs per chromosome, **(b)** Overview of chromosome 16 region enriched for maternal cluster mutations. X-axis and ideogram indicate chromosomal position. The red and blue histograms indicate the number of maternal cDNMs and paternal cDNMs identified in this study, respectively. The pale red and pale blue histograms indicate the number of maternal and paternal unclustered DNMs. The lowest track indicates normalized cSNP OG score, which is predictive for maternal DNMs. **(c)** The nucleotide substitution spectrum of maternal and paternal clusters and unclustered DNMs. Star indicates significance (pcO.OOl). **(d)** The nucleotide substitution spectrum of cDNMs by location, **(e)** The nucleotide substitution spectrum of polymorphism-derived clustered mutation by location. Star indicates significance (p<10^16^). (f) Region with increased maternal mutation rate on chromosome 2 (region displayed chrl6:l-100,000,000bp; region with maternal cDNMs chrl6:40,000,000-60,000,000).

To confirm these findings, we created a dataset of (unphased) clustered SNP variants based on publically available population-based genetic data^13^ **(Supplementary Methods).** This resulted in 1,146,891 clustered SNPs (cSNPs) across 522,487 clusters **(Supplementary Table 11).** We found that cSNPs on chromosomes that are enriched for maternal cDNMs are enriched for C>G substitutions (p<10^-16^, **Figure 2e, Supplementary Figure 8).** To further investigate this association, we calculated genome-wide score for C>G cSNP enrichment **(Supplementary Methods)** and found that the number of maternal cDNMs in a region is significantly correlated with high C>G scores (Poisson regression p<10^-16^ for maternal cDNMs, p=0.33 for paternal cDNMs, **Supplementary Figure 9).** Using this method we also identified an additional region on chromosome 2 that is enriched for maternal cDNMs **(Figure 2f).** This strong association between C>G scores of cSNPs with maternal cDNMs highlights maternal clusters' profound contribution to population polymorphisms in these regions.

The observed age-effect of maternal cDNMs suggests underlying mechanisms that are active during oocyte aging, a process that has been associated with the decreasing efficiency of double stranded break repair (DSBR)^14-16^. We therefore hypothesized that the maternal-aging associated clusters arise via a DSB-associated mechanism and investigated the occurrence of cDNMs at regions that are associated with DSBs. As proxies for DSB sites we used (1) sites of *de novo* meiotic gene conversion (MGC), (2) the flanking regions of *de novo* CNV breakpoints in our cohort, and (3) known recombination hotspots^17^.

We used MGC sites from Halldorsson et al.^18^ and found that these events co-localize with maternal cDNMs significantly more often than expected (p<0.004, bootstrapping, **Figure 3a, Supplementary Table 12).** This association is not significant for paternal MGCs with paternal cDNMs (p=0.056).

In our discovery cohort, we identified 45 high-quality *de novo* CNVs, of which 5 have a total of 17 DNMs within 100kb flanking the breakpoints **(Figure 3b, Supplementary Methods).** Exactly 15 of these 17 DNMs are cDNMs, which constitutes a high enrichment (p<2.2*10^-16^, Fisher’s exact test). For 6 of these 15 DNMs that the parent of origin was resolved, in all cases the DNMs arose from the maternal allele (p=0.03, Fisher’s exact test). In concordance with this, all 5 CNVs are deletions of the maternal allele **(Supplementary Table 13).** This association makes a single event as cause for both CNVs and cDNMs very likely. In our replication cohort, we also discovered 5 *de novo* deletion events. Two of these CNVs have a total of 4 DNMs from the same individual within 100kb of the CNV breakpoints, and two of these are within 20kb of each other **(Supplementary Figure 10),** again showing an enrichment of cDNMs (p=0.005, Fisher’s exact test). Interestingly, population CNV data showed a strong correlation between CNV breakpoints and cSNP density **(Figure 3c),** corroborated the co-segregation of CNV events and clustered mutations.

**Figure 3:**
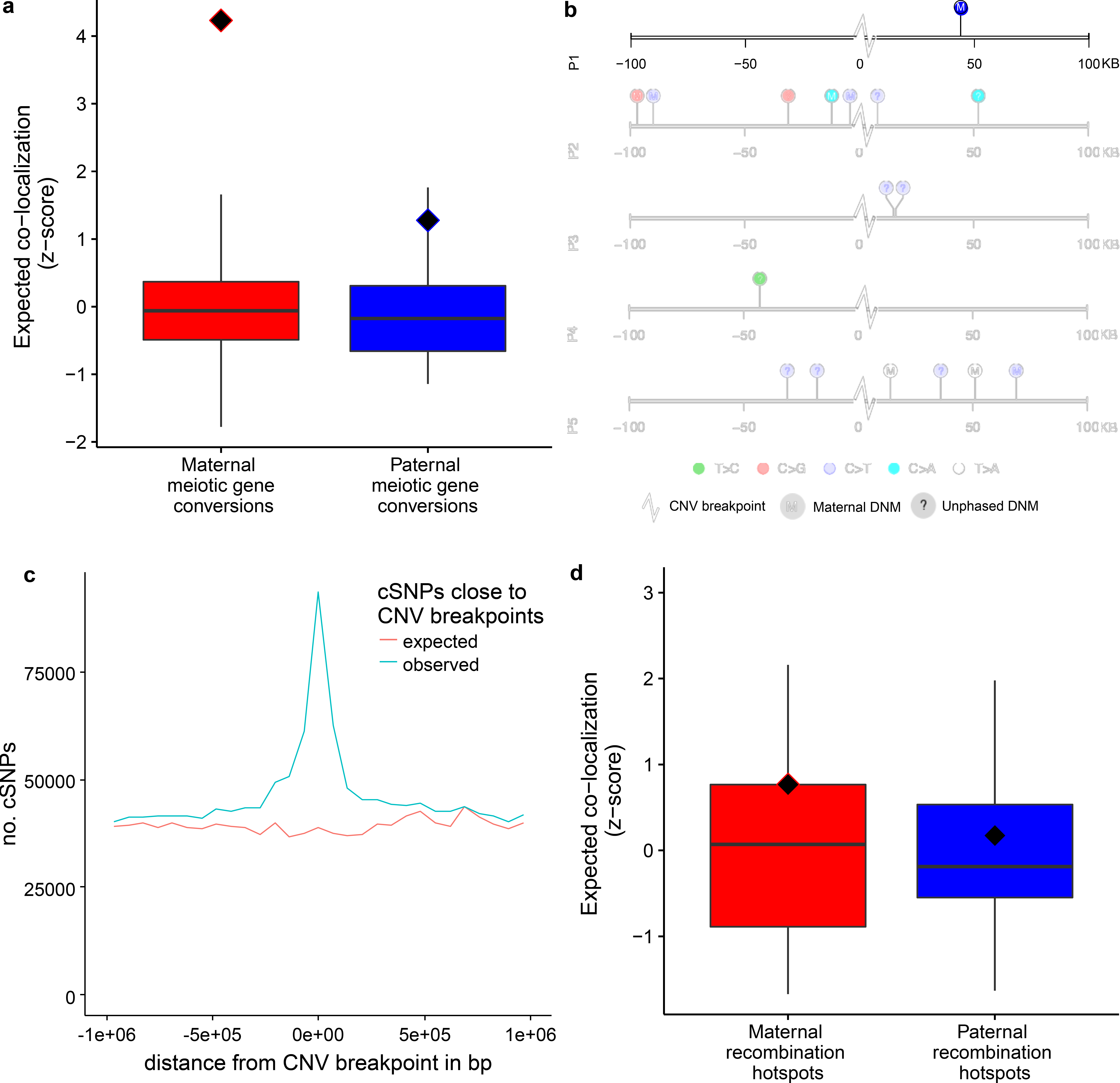
cDNMs and sites likely affected by DSBs. **(a)** Z-scores of expected and observed overlaps of cDNM clusters in our cohort and sex-matched meiotic gene conversion in another cohort. MCG: meiotic gene conversions, red diamonds: observed values, grey dots: simulated values, boxplot compartments: box: interquartile range; line: median; whiskers: extreme values <1.5 x interquartile ranges from box borders, **(b)** DNMs detected close to sites of *de novo* CNVs. **(c)** cSNP density close to CNV breakpoints, **(d)** Z-scores of expected and observed overlap cDNM clusters and sex-matched recombination hotspots. Symbols and boxplots as in (a).

Finally, we used gender specific recombination scores^17^ to assess whether cDNMs occur more often at regions of high recombination. Although we did not find a significant overlap of maternal cDNMs with regions of high maternal recombination (p=0.204 bootstrapping, **Figure 3d, Supplementary Table 12),** we did find that maternal recombination scores at maternal clusters are significantly higher than paternal recombination scores at paternal cluster locations (p=0.004, bootstrapping, **Supplementary Figure 11).**

The fact that the recombination rate increases with maternal, but not with paternal age, suggests that age-dependent cDNM accumulation in oocytes is due to abnormal recombination. Age-dependent accumulation of crossing-overs is likely to be the consequence from the repair of non-programmed DSBs^19^. In addition, during meiotic recombination, spatially proximal crossovers interfere with each other and observed escape from crossover interference is likely caused by non-programmed DSBs and therefore also increases with age. We found that that chromosomes 8, 9 and 16 are heavily enriched for maternal clusters and strikingly these chromosomes also have the highest degree of cross-over events escaping interference^19^. Finally, cDNM mutational spectra, and in particular those of maternal cDNMs, are very similar to the previously identified spectra of somatic mutations caused by deficiency in homologous recombination^20,21^ (Signature 3, **Supplementary Figure** 12), which is in agreement with a key role of non-programmed DSBs in the formation of maternal mutation clusters.

Overall our results show that abnormal recombination is likely to be the major force underlying the formation of clustered mutations. Future genome sequencing of larger families will allow us to identify individual crossing-over events and associate these with the presence of clustered mutations.

## URLs

goleft indexcov: https://github.com/brentp/goleft/tree/master/indexcov

## Acknowledgements

This study was funded by the Inova Health System with support from Fairfax County and the philanthropic support from the Odeen family. We thank the Inova translational medicine institute staff for supporting the study. We also thank the families who participated in the genomic studies that made this research possible. This work was partly financially supported by grants from the Netherlands Organization for Scientific Research (916-14-043 to C.G. and 918–15-667 to J.A.V.), and the European Research Council (ERC Starting grant DENOVO 281964 to J.A.V.).

## Author contributions

C.G., and J.E.N. designed the study. J.M.G., V.B.S., and W.S.W.W. performed the data analyses. W.S.W.W carried out QC, and *de novo* mutations calling. T.V. performed the Sanger validation. B.D.S., J.F.D, and J.E.N. supervised the data collection, sequencing and writing of the manuscript. D.B. assisted in data analyses and interpretation. J.M.G., V.B.S., W.S.W.W., J.A.V. and C.G. drafted the manuscript. All authors contributed to the final version of the paper.

## Online Methods

### Cohort

The cohort used in this study is from Inova Translational Medicine Institute’s Longitudinal Childhood Genome Study (previously referred to as the First 1,000 Days of Life and Beyond study), which represents a general population cohort not selected for health or disease^4^’^12^. The study was conducted by the Inova Translational Medicine Institute and approved by both the Inova and Western Institutional Review Boards (study 20120204). Parents and the newborns were recruited at Inova Fairfax Hospital between 2012 and 2014. A summary of participants' ages is given in **Supplementary Table** 1.

### Whole genome sequencing

Sample preparation, processing and whole-genome sequencing (WGS) have been previously described^4^’^12^. Briefly, DNA was extracted from peripheral blood obtained from each family member. Whole genome sequencing at ~40X was performed by Illumina Services (San Diego, CA) with the Illumina Whole Human Genome Sequencing Service Informatics Pipeline version 2.01-2.03. The sequenced reads were aligned to the hgl9 human reference genome by the ISAAC aligner^22^.

To systematically analyze the data quality of all sequencing reactions, a principal component analysis on summary statistics was performed **(Supplementary Figure 13).** The first principal component is highly correlated to average sequencing coverage; a group of outlying points refers to a group of sequencing reactions with average genome coverage above 70x. The second principal component is associated with the date of sequencing and the version of the software used for analysis, respectively. The third principal component is related to the estimated ancestries of the sequenced individuals.

### DNM Calling and Quality Control

Callable regions of each sample were determined by CallableLoci in GATK version 3.1. The number of callable bases by batch is shown in **Supplementary Figure 14.** Joint calling using HaplotypeCaller, PhaseByTransmission and ReadBackPhasing in GATK version 3.1 were performed on each of the 1,315 trios in the canonical autosomes. The putative de novo mutations were generated from taking PASS filter calls with heterozygous in the proband and homozygous reference in both parents in the PhaseByTransmission results in each trio. We have previously analyzed 816 trios^4^, of which, 65 trios were also sequenced by the Illumina services with pipeline version 2.0.0-2.0.1, and are not part of this cohort. These 65 trios sequenced by Illumina have gone through the same pipeline to generate a set of putative DNMs. We defined the positive set as those putative DNMs that overlap with previous identified DNMs identified using Complete Genomics (CG) technology (2,670), as well as those that were validated by Sanger sequencing (34), the total number in the true positive set is 2,704. The negative set consists of 50 random putative DNMs in each of the 65 trios that are not in the previously identified set by CG (50*65=3,250), as well as 4 false positive sites identified by Sanger, the total number of negative sites is 3,254. We note that some of the sites in the negative set are true positives but the number is likely to be low.

The test set which consists of the positive and negative sets was split by 90:10 ratio into training and test set. The R libraries randomForest version 4.6.10 and caret version 6.0.52 were used to train the random forest classifier. The OOB estimate of error rate on training set 1.77% and the error rate in the test set is 2.18%. The features used in the classifier and their relative importances are shown in **Supplementary Table 14.** The confusion matrix for the test set is shown in **Supplementary Table 15.**

In order to minimize the bias due to mapping errors and coverage differences, we further filtered the predicted DNMs by (1) callable regions in the cohort, which is defined by sequencing coverage was available for 90% of the samples, (2) good mappability regions, where mappable is defined according to the CRG lOOmer being equal to l^23^, sites also called by the Illumina Isaac pipeline, and sites with FS (FisherStrand test score) >=20, and sites with exceptionally high or low PL values **(Supplementary Table 16).** An overview of the filtering procedure is given in **Supplementary Table** 2.

In the initial sequencing cohort, there were 12 monozygotic twin pairs, 29 dizygotic twin pairs and a family of three trizygotic siblings. In order to assess the consistency in de novo calling, we investigated the concordance percentages of monozygotic and dizygotic families **(Supplementary Table** 17 and **Supplementary Table** 18). DNM calls in dizygotic twins are on average 95% concordant, the dizygotic average concordance is 0.1%. This is similar to concordance ratios observed previously^4^.

We removed 8 trios with large chromosomal anomaly in either the proband or one of the parents and removed (arbitrarily) monozygotic twin 2 in each set. After performing simple multiple linear regression, 4 samples have a significant Bonferroni p-value for studentized residuals (Bonferroni corrected p<0.05) and are removed from the cohort. We investigated the effect of average genome coverage on the filtered data. The results are shown in **Supplementary Figure 15.**

The method for determining the parent of origin of DNMs with Illumina WGS trio data was previously described^3^’^4^. Briefly, GATK PhaseByTransmission was used to assign parent of origin to informative heterozygous SNPs in the proband, GATK ReadBackPhasing was used to link DNMs to these informative SNPs. If contradictory markers are linked to the same DNM, it would not be assigned a parent-of-origin.

### Clustered DNMs

We defined cDNMs as DNMs on the same chromosome of the same individual within 20kb of each other. In order to estimate the chance of two DNMs being closer than 20kb on the same chromosome, we simulated 70,000 mutations at random positions within the callable and mappable genome. The randomized positions were given sample IDs as in the set of observed DNMs and the distances were calculated. We concluded that the false discovery rate of cluster detection is 0.045 at a threshold of 20kb **(Supplementary Table 4).** Statistics on the number of cDNMs per cluster are given in **Supplementary Table 3.**

### Clustered polymorphism variants

We use polymorphism data from the 1000 Genomes Project Consortium^13^. We only considered non-singleton variants with below 1% derived allele frequency, using the ancestral variant determined by The 1000 Genomes Project Consortium. Clusters were defined as two or more SNPs at distances between 10-1000 nucleotides from each other, such that all the genotypes carrying the derived allele for one of the SNPs also carry the derived allele for any other SNP within cluster. We show that cSNP spectra are similar to cDNM spectra: enriched by C>G mutations and depleted by CpG>TpG mutations, comparing with unclustered DNMs. We restricted ourselves to distance between cSNPs shorter than 1000 nucleotides, because spectra of larger clusters are progressively less similar to cDNMs. For analyzing the density of cSNPs around CNV breakpoints, the distance of cSNPs to CNV breakpoints on the same chromosome of the same haplotype (where available) were compared to the distances of cSNPs of random haplotypes to CNV breakpoints on the same chromosome.

### Statistical assessment of the maternal age effect

For analyzing the parental age effects on both the number of clusters as well as the number of cDNMs, linear models were fitted using the R statistical environment version 3.3.3. For comparing proband groups’ risks for having DNM clusters we used risk ratio statistics as implemented in the R package “epitools”. For assessing the enrichment of C>G substitutions on chromosomes 8, 9 and 16, we re-sampled the chromosome annotation 1000 times and compared the difference of the fractions of C>G mutations on the special chromosomes and the remaining autosomes to the observed value.

### Statistical assessment of DSB proxy regions overlap

For calculating distributions on the expected number of overlaps between DNM clusters and DSB proxy regions we used permutation testing as implemented in the R library Regione R^24^. DNM cluster regions were defined as the positions of cDNMs and the space between them. Recombination hotspots were defined as genomic sites with a recombination-score above 10^2^; Meiotic gene conversion sites were defined as the positions of conversed SNPs, the distances between them and a 20kb margin around them, as 20kb is the upper limit for inter-mutational distance within clusters. The cluster regions were randomized 500 times to genomic positions where at least 1000 base pairs were within the callable and mergable subset of the genome. For every randomization round the number of cluster positions overlapping DSB proxy regions was compared to the observed number of overlaps.

### De novo CNVs

In the discovery cohort, we called de novo CNVs using both coverage-based method FREEC^25^ and read-pair based method Manta^26^. We also calculated window based normalized coverage with “goleft indexcov”. For each proband, we called CNVs using the default options in FREEC with the proband as the case and one of the parents as control. We then required the CNVs subtracted from each parent to have 90% reciprocal overlap, with copy number equals 1 or 3, both parents have the mean normalized coverage between 0.85 and 1.15 in the region, the proband have mean normalized coverage smaller than 0.85 or greater than 1.15 in the region, with length greater or equal to 10kb. We performed joint calling for each trio with Manta using default options. We then filter for SV type being DEL or DUP, proband with GT equals to 0/1 and both parents with GT equal to 0/0, proband’s PR and SR for ALT allele >=3 and the proportion of PR and SR for ALT >=0.2, parents’ proportion of PR and SR for ALT <=0.05.

In the complete genomics data in the replication cohort, we required the de novo CNV to be called by both coverage based and read based methods. For the coverage based method, we first subtracted CNVs in the proband from one of the parents using the cnvSegmentsDiploidBeta files, and then we intersect the two putative de novo CNV files substracted from each parent, with 90% overlap, and size >9999. For the read based method, we subtracted highConfidenceSvEventsBeta file from the proband from allSvEventsBeta file from each of the parents, and intersected the two subtracted files requiring 90% overlap. The final list of de novo CNVs is generated by intersecting the coverage-based and read-based files from the same proband, requiring 90% overlap. Bedtools 2.22.0 was used to carry out region subtractions and intersections^27^.

### Mutation signatures

A large set of mutational signatures is known from cancer studies^20^, some of which are well annotated with mutational influences. To fit the patterns of our DNMs to these signatures we used an algorithm similar to the one described in^28^: a non-negative least-squares algorithm finds the mixture of known signatures that describes best the observed pattern. In order to get an indication of the robustness of the fitted mixture of signatures, we resampled DNMs from the original set with replacement and repeated the fitting procedure.

### Code availability

Code available upon request.

